# Bi-allelic variants in *TSPOAP1*, encoding the active zone protein RIMBP1, cause autosomal recessive dystonia

**DOI:** 10.1101/2020.05.24.086215

**Authors:** Niccolò E. Mencacci, Marisa M. Brockmann, Jinye Dai, Sander Pajusalu, Burcu Atasu, Paulina Gonzalez-Latapi, Christopher Patzke, Michael Schwake, Arianna Tucci, Alan Pittman, Javier Simon-Sanchez, Gemma L. Carvill, Bettina Balint, Sarah Wiethoff, Thomas T. Warner, Apostolos Papandreou, Audrey Soo, Reet Rein, Liis Kadastik-Eerme, Sanna Puusepp, Karit Reinson, Tiiu Tomberg, Joaquin Campos, Gabriela Pino, Hasmet Hanagasi, Thomas Gasser, Kailash P. Bhatia, Manju A. Kurian, Ebba Lohmann, Katrin Õunap, Christian Rosenmund, Thomas C. Südhof, Nicholas W. Wood, Dimitri Krainc, Claudio Acuna

## Abstract

Dystonia is a debilitating hyperkinetic movement disorder, frequently transmitted as a monogenic trait. Here, we describe homozygous frameshift, nonsense and missense variants in *TSPOAP1*, encoding the active zone RIM-binding protein 1 (RIMBP1), as a novel genetic cause of autosomal recessive dystonia in seven subjects from three unrelated families. Subjects carrying loss-of-function variants presented with juvenile-onset progressive generalized dystonia, associated with intellectual disability and cerebellar atrophy. Conversely, subjects carrying a pathogenic missense variant (p.Gly1808Ser) presented with isolated adult-onset focal dystonia. In mice, complete loss of RIMBP1, known to reduce neurotransmission, led to motor abnormalities reminiscent of dystonia, decreased Purkinje cell dendritic arborization, and reduced numbers of cerebellar synapses. In vitro analysis of the p.Gly1808Ser variant showed larger spike-evoked calcium transients and enhanced neurotransmission, suggesting that RIMBP1-linked dystonia can be caused by either reduced or enhanced rates of spike-evoked release in relevant neural networks. Our findings establish a direct link between presynaptic RIMBP1 dysfunction and dystonia and highlight the critical role played by well-balanced neurotransmission in motor control and disease pathogenesis.

## INTRODUCTION

Dystonia is a disabling movement disorder characterized by an excess of sustained or intermittent muscle contractions leading to abnormal, often repetitive, involuntary movements and postures (1). Dystonia, after Parkinson’s disease and essential tremor, is the third most common movement disorder and has an estimated frequency of 732 per 100,000 in the general population (2). Clinically, dystonia can occur as an isolated symptom or in combination with other movement disorders or neurological abnormalities (3).

Dystonia is thought to be a disorder of brain circuits involved in motor control,(4, 5) with several lines of evidence locating its origin in the basal ganglia(6) or in the cerebellar connections (7). The precise cellular and molecular events responsible for the genesis of dystonic movements are not understood, hindering the development of more effective treatments. Importantly, pathogenic variants in a growing number of genes have been causally linked to dominant and recessive Mendelian forms of dystonia (8), allowing the development of genetic animal models and the initial delineation of converging molecular pathways in disease mechanisms.(9) Along these lines, disruption of synaptic function has been recognized as a fundamental event in the pathogenesis of dystonia (10).

Synaptic function relies on fast and precise spike-triggered neurotransmitter release at the presynaptic active zone (11). The active zone is a cytomatrix composed of several large protein families, including Munc13s, RIMs, ELKs, α-liprins, and RIM-binding proteins (12). These proteins contain multiple protein-protein interaction domains, which facilitates the formation of a densely interconnected protein network that enables docking and priming of synaptic vesicles for exocytosis and ensures rapid and efficient coupling between presynaptic calcium (Ca^2+^) influx and synaptic vesicle fusion (13).

RIM-binding proteins (RIMBPs) are central components of the active zone that, in conjunction with RIMs, determine the precise localization of presynaptic voltage-gated Ca^2+^ channels (VGCCs) and ensure tight coupling between presynaptic action potentials and synaptic vesicle exocytosis (14, 15). RIMBPs comprise two main brain isoforms, RIMBP1 and RIMBP2, which are structurally similar and consist of a SH3 domain at the N-terminal region, followed by three central fibronectin type-3 domains (FN3), and two additional SH3 domains in their most distal C-terminal regions.(16) Mechanistically, it is *via* their SH3 domains that RIMBPs bind to the proline-rich sequences in the cytosolic domain of presynaptic VGCCs, tethering them to the active zone (14, 17), and controlling the precision and fidelity of neurotransmitter release and synaptic transmission (18, 19).

Here, we demonstrate that rare homozygous truncating and missense variants in *TSPOAP1* (MIM* 610764; previously known as *BZRAP1*), the gene encoding RIMBP1, are causally associated with autosomal recessive dystonia in three unrelated families. In mice, deletion of RIMBP1 caused motor abnormalities reminiscent of dystonia as well as changes in the biochemical composition and morphology of the cerebellum. Interestingly, the pathogenic missense variant rendered abnormally large presynaptic Ca^2+^ entry and increased transmitter release in response to presynaptic firing. Altogether, our results highlight the critical role played by precisely fine-tuned synaptic transmission in normal motor function, and establish a direct link between RIMBP1- mediated neurotransmitter vesicle release and dystonia pathogenesis.

## Results

### Homozygous TSPOAP1/RIMBP1 variants cause autosomal recessive dystonia

Using a combination of homozygosity mapping and whole-exome sequencing (WES), homozygous variants in *TSPOAP1* were independently identified as the top candidate genetic cause of autosomal recessive dystonia in the three families reported herein. The online platform GeneMatcher (20) was used to share results among the three groups that identified the variants.

**Family A** is a consanguineous pedigree of Gujarati Indian origin (parents of the affected subjects are first cousins) (Figure 1A, left). Homozygosity mapping revealed a single candidate chromosomal region corresponding to a large run of homozygosity (ROH) of ∼11,5 Mb on chromosome 17 (chr17:53,415,850–64,831,657), which was unique to the three affected family members. After applying the filtering strategy to WES data, only a single candidate causative variant remained, a frameshift deletion in *TSPOAP1* (NM_004758.3:c.538delG, p.Ala180Profs*8) located within the disease-associated ROH and shared by all three affected subjects. Sanger sequencing confirmed the presence of the variant and showed that both parents were heterozygous carriers while the healthy sibling was homozygous for the reference allele, thus confirming complete disease co-segregation. This frameshift variant is absent in more than 120,000 subjects listed in gnomAD, which contains >15,000 subjects of South Asian ancestry.

**Figure 1.**
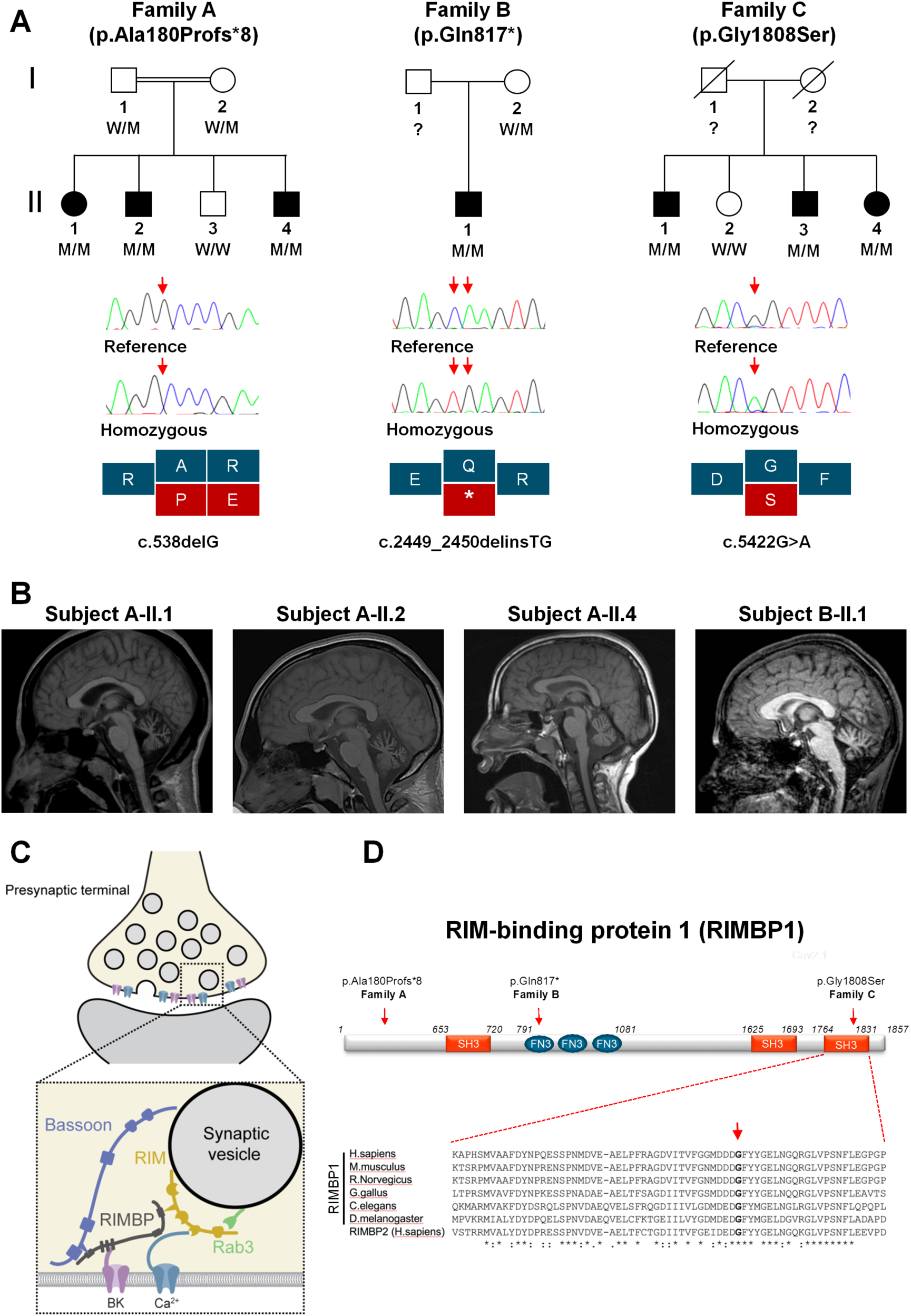
Pedigrees, genetic findings, and radiological features of subjects with pathogenic *TSPOAP1/*RIMBP1 variants. **A.** Top. Pedigrees and variant status of affected (closed symbols) and healthy (open symbols) members of the three families with bi-allelic *TSPOAP1* pathogenic variants. Bottom. Sanger sequencing validation of the variants and schematic representation of the variant effect on protein coding sequence. The following abbreviations are used: WT for wild-type alleles and M for mutant alleles. **B.** Brain MRI sagittal images T1-weighted images of subjects A-II.1 (age 28), A-II.2 (age 23), A-II.4 (age 13) and B-II.1 (age 17), demonstrating cerebellar atrophy with a predominance of the vermis in all four subjects carrying homozygous loss-of-function variants. **C.** Cartoon illustrating the molecular interactions of RIM-binding proteins (RIMBPs) at the presynaptic active zone. RIMBPs bind with the first SH3 domain to Bassoon, with the FN3 domains to calcium-activated potassium channels (BK) and with the second and third SH3 domain to voltage-gated calcium-channels (Ca^2+^) and RIM proteins. **D.** Schematic representation of RIMBP1 structure including protein domains and localization of the identified variants. The amino acid residue p.Gly1808 is located in the C-terminal SH3 domain and shows complete evolutionary conservation across all species and in the human protein homolog RIMBP2. Asterisks indicate invariant residues (full conservation), whereas colons and periods represent strong and moderate similarities, respectively.

**Family B** is of Estonian origin and has been previously described.(21) There was no reported history of parental consanguinity. The proband is the only child in the family (Figure 1A, middle). Chromosomal microarray analysis did not detect CNVs. Homozygosity mapping revealed three large ROHs (chr4: 10,769,440 - 21,449,134; chr10:54,942,171-63,972,930; chr17:55,928,909-67,419,074) indicating likely unreported consanguinity. WES analysis identified a homozygous STOP-gain variant in *TSPOAP1* (NM_004758.3:c.2449_2450delinsTG, p.Gln817*), located in the largest ROH as the best candidate. Sanger sequencing confirmed the presence of the homozygous variant in the proband and heterozygous in the mother (father’s DNA was not available). This variant is not present in gnomAD, which contains 2,418 Estonian subjects.

**Family C** is of Turkish origin (Figure 1A, right). There was no reported consanguinity in the family but the parents came from the same village. Three large ROHs (chr8: 33,406,218 - 42,407,799; chr11: 56,184,803 - 82,708,284; and chr17:54,967,949 - 66,426,255) were found exclusively in the proband. WES analysis revealed three rare homozygous coding variants located within the identified ROHs, *KCNU1* (NM_001031836: c.1280T>C, p.Ile427Thr [rs570237282]), *NADSYN1* (NM_018161:c.278G>A, p.Arg93Gln [rs763061270]) and *TSPOAP1* (NM_004758.3:c.5422G>A, p.Gly1808Ser [rs752560074]). Amongst these, only the variant in *TSPOAP1* was homozygous also in the other siblings affected by dystonia. The variant is predicted pathogenic by all *in silico* prediction tools and has a CADD score of 27.7. The variant is not present in the homozygous state in gnomAD (minor allele frequency 0.00001061) and is not present in the Great Middle East Variome database (22). Finally, the variant was absent in a further ∼1000 Turkish families with unrelated medical conditions, thus excluding it as a common variant specific to the Turkish population.

*TSPOAP1* variants identified in families A and B are both predicted to render truncated RIMBP1 products lacking most if not all functional domains. Thus, the carriers of these variants represent essentially RIMBP1 knockout (KO) subjects (Figure 1D). In contrast, the p.Gly1808Ser variant found in Family C is a missense variant located in the last RIMBP1 SH3 domain, which is critical for its binding to VGCCs (14, 17), and shows complete evolutionary conservation across species down to invertebrates and in the human homolog RIMBP2 (Figure 1D).

Loss-of-function variants are absent in ∼ 15,000 in-house exomes sequenced at the three centers who analyzed the families and homozygous loss-of-function variants are not seen in a further ∼130,000 subjects listed on gnomAD. Furthermore, we did not observe any homozygous rare *TSPOAP1* missense variant located within any of the three functionally relevant SH3 domains in both in-house exomes and gnomAD, indicating that homozygous variants in these regions are exceedingly rare and adding further evidence in support of the pathogenic role of the variant identified in family C. Finally, mining WES data from an additional ∼700 cases affected with different forms of dystonia, failed to identify additional cases with confirmed bi-allelic *TSPOAP1* changes.

### Clinical features of patients with bi-allelic TSPOAP1/RIMBP1 variants

The clinical features of all subjects with *TSPOAP1* variants are summarized in Table 1. In brief, all four subjects with homozygous loss-of-function *TSPOAP1* variants (Family A and B) shared a strikingly similar phenotype characterized by normal motor and language development, mild learning disabilities that first appeared in primary school, and onset in early teenage years of progressive generalized dystonia (Supplemental video 1). Dystonia displayed a clear craniocaudal gradient in all four subjects, with dystonic movements severely affecting the cranio-cervical district and to a lesser extent the trunk and the four limbs. The impairment of the cranial muscles was particularly severe, resulting in prominent dysphonia and dysarthria and progressive difficulties with swallowing. Upper limb examination showed continuous dystonic posturing and writhing movements, resulting in impaired dexterity and severe handwriting difficulties. Gait examination showed progressive abnormalities including bilateral foot inward turning, tiptoeing, leg stiffening, and abnormal knee flexion, resulting in progressive loss of autonomous ambulation.

**Table 1.** Clinical features of subjects with homozygous pathogenic *TSPOAP1* variants.

|  | A-II.1 | A-II.2 | A-II.4 | B-II.1 | C-II.1 | C-II.3 | C-II.4 |
| --- | --- | --- | --- | --- | --- | --- | --- |
| **Gender** | Female | Male | Male | Male | Male | Male | Female |
| **Origin** | Indian Gujarati | Indian Gujarati | Indian Gujarati | Estonian | Turkish | Turkish | Turkish |
| **Age at last examination** | 31 | 26 | 20 | 23 | 73 | 73 | 68 |
| ***TSPOAP1* variant<sup>a</sup>** | c.538delG, p.Ala180Profs*8 | c.538delG, p.Ala180Profs*8 | c.538delG, p.Ala180Profs*8 | c.2449_2450delinsTG, p.Gln817* | c.5422G>A, p.Gly1808Ser | c.5422G>A, p.Gly1808Ser | c.5422G>A, p.Gly1808Ser |
| **Motor development** | Normal | Normal | Normal | Normal | Normal | Normal | Normal |
| **Dystonia characteristics** |  |  |  |  |  |  |  |
| **Age at onset of motor symptoms (y)** | 12 | 12 | 13 | 11 | ~60s | 67 | 58 |
| **Symptoms at presentation** | Loss of voice and speech difficulties | Speech difficulties | Rapid onset of speech difficulties | Dystonic movements in right hand and leg | Hand tremor | Cervical dystonia | Cervical dystonia |
| **Symptoms of cranial, cervical, laryngeal dystonia (onset, y)** | Severe orofacial involvement with opening spasms, severe dysarthria and dysphonia, dysphagia, severe torticollis and retrocollis (12) | Severe laryngeal dystonia (whispering dysphonia), dysarthria (12) | Anarthria and severe oral dystonia, severe dysphagia, frequent spasms of platysma muscle activated by attempts to speak (13) | Mild dysarthria progressed into severe dysarthria and orofacial-mandibular dystonia, blepharospasm, retrocollis and laterocollis (14) | None | Irregular dystonic horizontal head tremor, mild right torticollis | Retrocollis, right laterocollis, mild left torticollis, right shoulder elevation |
| **Upper limb involvement (onset, y)** | Continuous dystonic posturing and movements worsened by action, muscle overflow, impaired dexterity and handwriting difficulties (17) | Continuous dystonic posturing and movements worsened by action, muscle overflow, impaired dexterity and handwriting difficulties (16) | Continuous dystonic posturing and movements worsened by action and with muscle overflow, impaired dexterity and handwriting difficulties (14) | Continuous dystonic posturing and movements worsened by action, muscle overflow, impaired dexterity and handwriting difficulties (11) | Dystonic hand resting tremor | Bilateral postural tremor | Mild bilateral writhing movements |
| **Onset of lower limb involvement (onset, y)** | Tiptoe walking with extreme knee flexion (17) | Bilateral foot inward turning (16) | Bilateral foot inward turning (16) | Bilateral foot inward turning (11) | None | None | None |
| **Distribution at final examination** | Generalized with cranio>caudal gradient | Generalized with cranio>caudal gradient | Generalized with cranio>caudal gradient | Generalized with cranio>caudal gradient | Focal | Focal | Focal |
| **Gait** | Able to walk only with support; severe dystonic features | Possible only for short distances; dystonic gait | Possible only for short distances; bilateral leg dystonic posturing, narrow-based | Possible only for short distances, very unsteady, tiptoe walking, knee flexion and severe dystonic posturing, narrow-based | Normal | Normal | Normal |
| **Cognition** | Learning disability since school; neuropsychology testing at age 27 revealed extensive cognitive impairment | Learning disability since school; cognitive decline (not formally tested) | Learning disability since school; cognitive decline (not formally tested) | Progression from mild (11y) to moderate intellectual disability (16y) | Mild cognitive impairment | Mild cognitive impairment | Mild cognitive impairment |
| **Cerebellar signs** | None | Square wave jerks, slow and hypometric saccades, mild intentional tremor and dysmetria in 4 limbs, broad base | Square wave jerks, slow and hypometric saccades, mild dysidiadocokinesia in UEs | None | None | None | None |
| **Other neurological features** | None | Episodes triggered by sudden movements, featuring painful truncal spasms, responsive to AEDs but with no EEG correlate. | Idiopathic generalized epilepsy with generalized tonic-conic seizures with onset at age 4, resolved by age 10 | Rigidity in right limbs | None | None | Bilateral Babinski |
| **Brain MRI** | Cerebellar atrophy | Cerebellar atrophy | Cerebellar atrophy | Cerebellar atrophy | NA | NA | Reported as normal (Images NA) |
| **Treatment trials** | BCF, TBZ, LD, BT (not effective). Currently on CNZ 0.5 mg BID and THP 2 mg OD | LD (no benefit). Currently on THP 2mg (minimal improvement) and PHT | THP, BT (not effective), currently on CNZ | LD, CBZ (no benefit). Currently on LV (some initial benefit), TBZ (some benefit), BT (some benefit) | None | None | THP, BCF and BT injections (beneficial) |
| **Previous negative genetic testing** | *TOR1A*, *THAP1*, common mitochondrial DNA mutations | *SCA1*, *SCA2*, *SCA3*, *SCA6*, *SCA7*, *SCA12*, *SCA17*, *DRPLA*, *CSTB*, *FTXN*, *TOR1A* | *TOR1A*, *GCH1*, *SGCE*, *ATM*; normal chromosomal microarray | *TH*, *GCH1*, *HTT*, *ATP7B* | None | None | *HTT* |
| **Previous investigations** | Normal copper and ceruloplasmin, ferritin, acanthocytes, serum and urine amino and organic acid, VLCFA, blood smear, phytanic and pristanic acid | Normal white blood cell enzyme analysis, purine metabolism, CSF (except marginal 5-MTHF deficiency) | Normal EMG and NCS | Normal serum and urine amino and organic acids, MPS and sialic acid, purine/pyrimidines, CSF (except diminished level of 5-MTHF). Muscle biopsy (scattered angular atrophic muscle fibers referring to denervated fibers) | None | None | Normal EMG and NCS |

AEs=antiepileptic drugs, BCF=baclofen, TBZ=tetrabenazine, LD=levodopa, BT=botulinum toxin, CNZ=clonazepam, THP=trihexyphenidyl, PHT=phenytoin CBZ=carbamazepine, LV=leucovorin, EMG=electromyography, NCS= nerve conduction studies. 5-MTHF=5-methyltetrahydrofolate. MPS= mucopolysaccharides NA=not available <sup>a</sup> NCBI transcript NM\_004758.

All four cases showed clear evidence of cognitive deterioration. Formal cognitive assessment was performed in case A-II.1 (age 27; extensive cognitive impairment) and in subject B-II.1 (age 10 and 17; progression from mild to moderate intellectual disability). Other additional neurological features that were only variably observed included generalized tonic-clonic seizures (subject A-II.4; EEG showed bilateral interictal epileptic activity), superimposed episodes of limb and painful truncal dystonia triggered by sudden movements and controlled by treatment with phenytoin (subject A-II.2), and lower limb spasticity with hyperactive reflexes (subject B-II.1).

Subjects from family C carrying the homozygous missense change p.Gly1808Ser presented with a milder dystonic phenotype, characterized by adult-onset focal dystonia or dystonic tremor (supplemental video 2). The index proband (C-II.4) developed dystonia at age 58 and on examination she had segmental dystonia affecting her neck and upper limbs. Two of the proband’s siblings presented in their 60s with either isolated tremulous cervical dystonia and upper limb postural tremor (subject C-II.3) or isolated hand dystonic tremor (subject C-II.1).

### Patients with homozygous TSPOAP1/RIMBP1 loss-of-function variants display progressive cerebellar atrophy

Surprisingly, brain MRI showed prominent cerebellar atrophy with a predominance for the vermis in all four subjects from Families A and B (Figure 1B), a finding that is highly atypical in cases with generalized dystonia. Serial brain imaging in subject A-IV.2 (MRI at age 5, 8 and 14) and B-II.1 (MRI at age 15 and 17) documented progression of the cerebellar atrophy. The prominence of this radiological finding was in striking contrast with the paucity of classic clinical features of cerebellar dysfunction (i.e. ocular nystagmus, gait ataxia, limb dysmetria). Conversely, supratentorial, brainstem and spinal cord parenchyma had otherwise a normal appearance and there were no signal abnormalities or volume loss in the basal ganglia. Brain imaging was reported as normal in the proband of family C, but images were not available for direct review.

Brain expression profile of *TSPOAP1/*RIMBP1 and its homolog *RIMPB2* supports an essential role for RIMBP1 in cerebellar Purkinje neurons. Previous studies (16) as well as gene expression datasets from human and mouse brains, including GTEx (23) (human brain expression) and Brain Allen Atlas (mouse brain expression) (24), showed that both *TSPOAP1* and its homolog *RIMBP2* are highly expressed across several brain regions in a partly overlapping and partly segregated manner (Suppl. Figure 1). Importantly, in the cerebellum the expression of *TSPOAP1/*RIMBP1 was very high, particularly in Purkinje cells, while the expression of *RIMBP2* was low throughout. A similar pattern of expression was observed in the striatum as well, though less pronounced than in the cerebellum.

RIMBP1 and RIMBP2 are functionally redundant and can compensate for each other’s function when individually deleted (18, 19). The particular RIMBP expression pattern in Purkinje cells indicates that compensation might not occur in these cells, and thus suggests Purkinje cells might be particularly susceptible to RIMBP1 dysfunction caused by the identified *TSPOAP1* pathogenic variants.

### Genetic deletion of RIMBP1, but not RIMBP2, triggers motor abnormalities in mice

To support the pathogenic role of the identified homozygous loss-of-function RIMPB1 variants, we pursued further studies in mice. For this, we generated RIMBP1-KO mice, as previously reported (25) (Supp. Figure 2A). RIMBP1-KO mice displayed normal weight and no obvious developmental abnormalities (Supp. Figure 2B), indicating that constitutive removal of RIMBP1 has no significant effects on mouse survival and gross development.

We then tested whether RIMBP1-KO impacts mouse motor behaviors that are commonly disrupted in dystonia models. First, quantification of open field activity showed that KO mice displayed ∼30% overall increase in spontaneous locomotor activity compared to WT littermates (Figure 2A). Second, in a beam-walk motor coordination test (Figure 2B), RIMBP1-KO mice showed nearly 100% more ‘slips’ and were significantly slower at crossing the beam than WT littermates (Supplemental video 3). Third, suspension of mice by their tail (Figure 2C and Supplemental video 4) for 10 or 20 seconds led to limb-clasping behavior in 80% of RIMBP1-KO mice and this percentage increased to 100% when the suspension lasted for 30 seconds (averaged clasping index of ∼1.5±2.5). In striking contrast, none of the WT mice exhibited hindlimb-clasping in any of these conditions (average clasping index ∼0.5±0.1 Figure 2C, middle). Fourth, on rotarod testing, we did not detect significant differences between RIMBP1-KO and WT in their initial coordination nor in their learning rate (Supp. Figure 2C).

**Figure 2.**
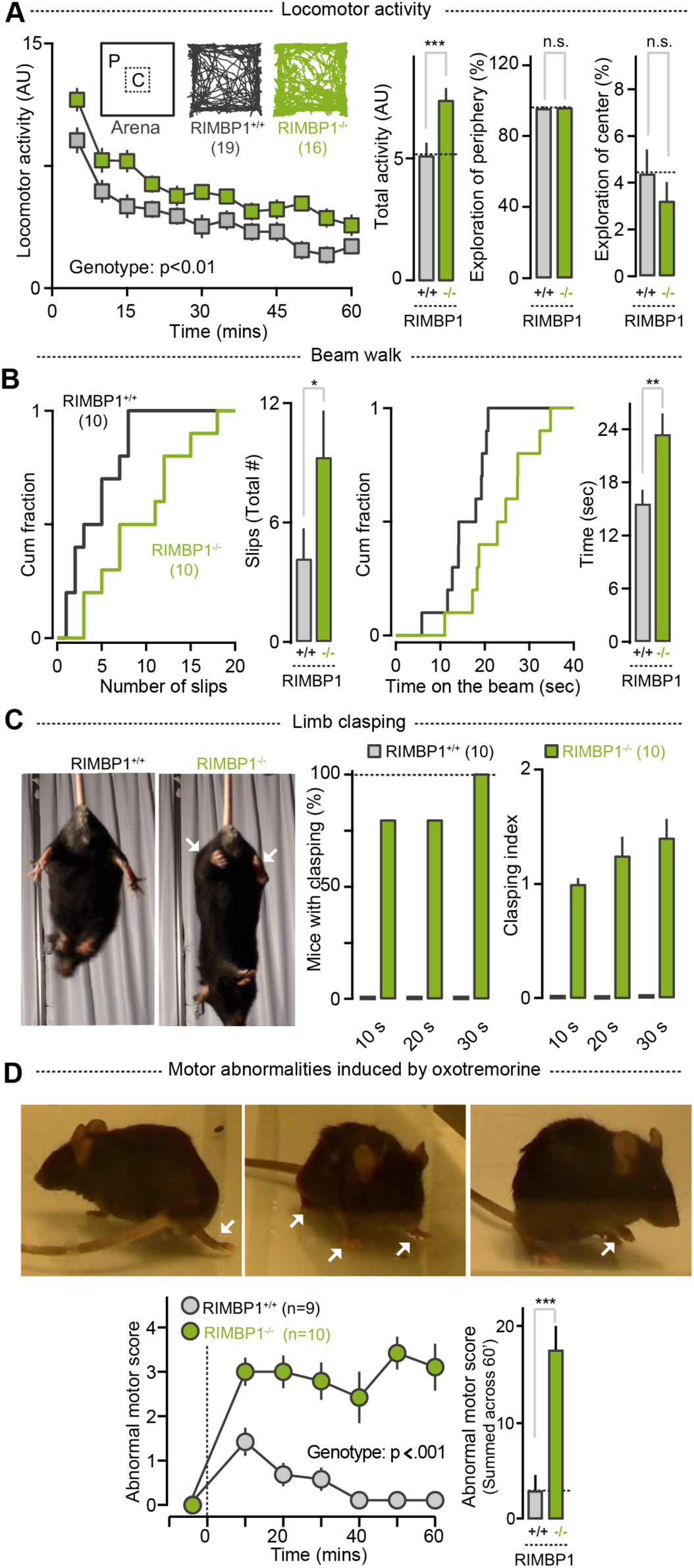
Motor abnormalities in RIMBP1-KO mice. **A.** Open-field locomotor activity. Left. Locomotor activity as a function of time (bin width: 5 mins). Middle. Total activity in RIMBP1-WT and RIMBP1-KO mice. Right. Comparison of relative activity in the periphery (P, left) and the center (C, right) of the open field arena in control and mutant mice. **B.** Beam-walk test. Left. Cumulative distribution (left) and summary graphs (right) of total time to cross the beam in RIMBP1 WT and RIMBP1-KO mice. Right. Cumulative distribution (left) and summary graphs (right) of total number of slips in RIMBP1 WT and RIMBP1 KO mice. **C.** Limb clasping test. Left. Representative pictures of a RIMBP1 WT (left) and a RIMBP1 KO (right) during the tail suspension test used to measure limb clasping. Arrows highlight limb clasping in the RIMBP1 KO mouse. Left. Percentage of RIMBP1-WT and RIMBP1-KO mice displaying limb clasping in tail suspension test lasting 10, 20, or 30 secs. Right. Clasping index in control and mutant mice tail-suspended for 10, 20, or 30 secs. **D.** Abnormal movements and postures triggered by systemic oxotremorine. Top. Examples of abnormal postures (arrows) found in RIMBP1-KO after oxotremorine treatment (0.01mg/Kg). Bottom, Left. Time course of abnormal motor scores after oxotremorine treatment (0.01mg/Kg). Motor scores were assessed in 2 minutes bins. Abnormal motor score scale: 0=Normal motor behavior; abnormal motor scores scale: 1= No impairment but slightly slowed movements; 2= Mild impairment: occasional abnormal postures and movements; ambulation with slow walk; 3=Moderate impairment: frequent abnormal postures and movements with limited ambulation; 4=Severe impairment: sustained abnormal postures without any ambulation or upright position. Bottom, right. Summed abnormal motor scores recorded for 60 minutes after drug injection. Number of experiments: A, right: 19 WT, 16 KO; B: 10 WT, 10 KO; C: 10 WT, 10 KO; D: 9 WT, 10 KO.

As human and mouse brains display high RIMBP1 and low RIMBP2 expression in the striatum and the cerebellum, we hypothesized that selective lack of RIMBP1 but not of RIMBP2 would lead to motor abnormalities. Consistent with the expectation, and in striking contrast to RIMBP1-KO, RIMBP2-KO neither affected the overall locomotor activity or beam walk performance, nor caused limb-clasping behavior (Suppl. Figure 3C-G). Thus, deletion of RIMBP1 but not RIMBP2 leads to motor abnormalities in mice.

### Abnormal movements and postures in RIMBP1-KO mice upon acetylcholine muscarinic receptor stimulation

While RIMBP1-KO mice displayed several baseline motor abnormalities, we did not observe any spontaneous abnormal movements or postures, the clinical hallmark of dystonia. Therefore, we challenged RIMBP1-KO mice with the non-selective muscarinic receptor agonist oxotremorine, which has been recently shown to promote overt dystonia-like abnormal movements and postures in another genetic mouse model of dystonia (26). RIMBP1-WT and -KO mice were injected with low doses of oxotremorine (0.01 mg/kg intraperitoneally [i.p.]), and then the appearance of abnormal movements and postures was assessed for up to 1-hour post injection (Figure 2D).

RIMBP1-WT mice displayed only mildly reduced spontaneous locomotion and rare events of head and body tremors or hindlimb extension during the first 30 minutes post injection only. In striking contrast, littermates RIMBP1-KO mice showed prominent and prolonged abnormalities in postures and movements that lasted for at least 1-hour post oxotremorine injections (Figure 2D, Bottom, left; supplemental video 5). These included a dramatic and sustained reduction in spontaneous locomotion, prolonged periods of immobility with hunched trunk postures associated with flexion of hindlimbs and extension of forelimbs, slow and uncoordinated gait with extended body and abnormal hindlimb postures, and frequent excessive jerking of the head and body twitching, prolonged motionless standing on hindlimbs with uncoordinated movements of the forelimbs and occasional raising of a single paw (Suppl. Video 6). A quantitative analysis, using an established scale for motor abnormalities in mice (26), estimated that oxotremorine increased motor dysfunction in RIMBP1-KO by about 300% compared to WT littermates (Figure 2D, Bottom, right), indicating a severe susceptibility to muscarinic cholinergic stimulation caused by RIMBP1 deletion.

### Cerebellar morphology and protein composition in mice lacking RIMBP1

Given the presence of significant cerebellar atrophy in subjects with homozygous RIMBP1 truncation and the high expression levels of RIMBP1 in cerebellar Purkinje cells, we asked whether RIMBP1 deletion is causally linked to morphological or biochemical abnormalities in the mouse cerebellum. Gross visual inspection of cerebelli from RIMBP1-WT and KO mice did not show obvious size differences (Figure 3A, left) and quantitative analyses did not detect a significant difference in the cerebellar volume (Figure 3A, left) or the total number of Purkinje cells (Figure 3A, right). Furthermore, immunoblotting analysis of cerebellar lysates did not show significant changes in the levels of general neuronal markers between RIMBP1-WT and KO mice (Suppl. Figure 3).

**Figure 3.**
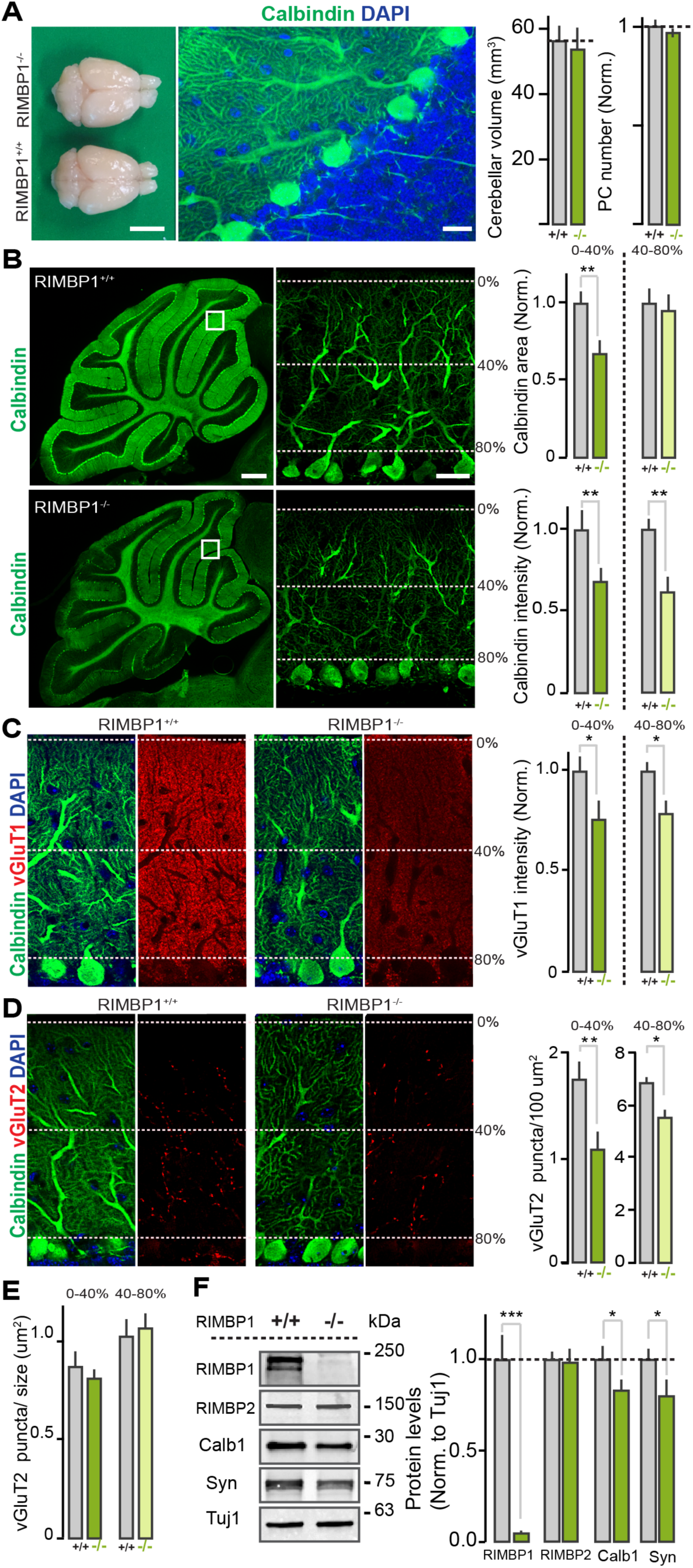
Cerebellar abnormalities in RIMBP1-KO mice. **A**. Gross anatomy of the cerebellum. Left. Dorsal view of brains from a RIMBP1-WT and a RIMBP1-KO mouse. Middle. Confocal image of a parasagittal cerebellar section stain with DAPI and antibodies anti-calbindin (green) to label and count Purkinje cells. Right. Summary plot of cerebellar volume in RIMBP1 control and mutant mice (left), and of Purkinje cells number quantified in parasagittal cerebellar sections (right). **B**. Structure of Purkinje cell dendrites. Left. Representative low-magnification parasagittal cerebellar sections stained with anti-Calbindin antibodies in a RIMBP1-WT (top) and a RIMBP1-KO mouse (bottom). Middle. High-magnification confocal images (single optical sections) of the areas indicated with the white boxes on the left displaying outer (0-40%) and inner (40-80%) Purkinje cell dendritic domains. Right. Top, summary graphs of calbindin coverage (area) in different Purkinje cell dendritic domains in RIMBP1 control and mutant mice. Bottom, summary graphs of calbindin intensity signal in the same PC dendritic domains. **C**. Organization of parallel fiber synapses. Left, representative confocal images of cerebellar sections from a RIMBP1-WT mouse. Middle, representative images from a littermate RIMBP1-KO mouse. Sections were stained for calbindin (green, to label Purkinje cells), for vGluT1 (red, t0 visualize parallel fiber inputs), and for DAPI (blue, cell nuclei). Right, summary graphs of vGluT1 intensity for the outermost and innermost Purkinje cell dendritic domains in RIMBP1 WT and littermates KO mice. **D, E**. Same as in C but for climbing fiber synapses stained with antibodies anti-vGluT2. In this case the density (**D**, right) and the size (**E**) of vGluT2-containing puncta in RIMBP1-WT and littermates KO mice, was compared. **F.** Levels of key proteins in cerebellar lysates from RIMBP1-WT and RIMBP1-KO mice. Left, representative blots. Right. Summary plots of band intensity values normalized to Tuj1. Number of experiments (mice/sections): **A**. Left: 5 WT, 6 KO. Right: 6/62WT, 6/47 KO**. B**. Intensity: 6/59 WT, 6/64 KO. Area: 6/57 WT, 6/60 KO**. C**. vGluT1: 6/59 WT, 6/56 KO. **D, E**. vGluT2: 6/54 WT, 6/60 KO.

We found, however, that RIMBP1-KO led to significant morphological abnormalities in the dendritic arbors of Purkinje cells (Figure 3B). Quantification of the integrity of Purkinje cell, assessed by measuring the total area covered by Calbindin immunolabeling in parasagittal cerebellar sections, revealed a ∼30% loss of coverage in the more distal portion of Purkinje cell dendritic arbors (Figure 3, right). Furthermore, the overall Calbindin intensity was significantly decreased throughout the cerebellum (31.6% reduction in the distal domain; 38.4% in the proximal domain, Figure 3B, right). This result was confirmed by the immunoblotting from cerebellar lysates, which showed a ∼20% reduction of the Calbindin signal in RIMBP1-KO mice (Figure 3D).

We also found that RIMBP1 deletion led to changes in the number of excitatory but not inhibitory synapses onto Purkinje cells. Purkinje cells receive two major excitatory inputs, parallel fibers and climbing fibers, which can be readily identified by expression of vesicular glutamate transporters vGluT1 or vGluT2, respectively. RIMBP1 deletion reduced both proximal and distal dendritic vGluT1 staining intensity by ∼20% compared to WT controls (Figure 3C), whereas vGlut2-containing clusters were reduced by around ∼30% in the distal and by ∼15% in the proximal dendritic arbors (Figure 3D, E). Again, these results were confirmed by measurements of synapsin levels by immunoblotting from cerebellar lysates, which showed a ∼20% reduction in RIMBP1-KO mice (Figure 3F).

Finally, immunoblotting analysis showed that neither the level of several active zone proteins (including RIM1 and RIMBP2) (Figure 3D, and Suppl. Figure 4) nor the levels of the SNAREs and associated proteins were significantly different in WT and KO cerebelli. Likewise, the levels of VGCCs and BK-channels that interact with RIMBPs, as well as the levels of several postsynaptic proteins found in excitatory and inhibitory synapses, were not altered (Suppl. Figure 4).

Together, these results indicate that constitutive deletion of RIMBP1 does not affect cerebellar volume, biochemical composition, and the number of Purkinje cells, but significantly impacts Purkinje cell calbindin levels, dendritic morphology, and the number of excitatory synaptic inputs from both climbing and parallel fibers onto these cells.

### Impact of the pathogenic RIMBP1 p.G1808S substitution on synaptic transmission

Subjects from family A and B described above carry *TSPOAP1* truncating variants that essentially lead to the loss of all RIMBP1 functional domains. Therefore, these variants are expected to cause synaptic dysfunctions reminiscent of those observed in RIMBPs KO neurons, including reduced priming, uncoupling of VGCCs from the active zone, and reduced synaptic transmission fidelity (18, 19, 27, 28).

Subjects from family C, in contrast, carry a missense variant (p.Gly1808Ser) that changes a single amino acid in the last SH3 domain of RIMBP1. Patients carrying this variant displayed a different dystonic phenotype, suggesting that the pathogenic mechanism triggered by this variant may be different from that of the truncating RIMBP1 variants. To address this, we used autaptic neuronal cultures (29), which allow for precise assessment of synaptic parameters (30) (Figure 4A). Because of the known functional redundancy between RIMBPs and RIMs,(19) we performed all experiments in neuronal cultures prepared from RIMBP1,2/RIM1,2 quadruple conditional KO mice (qKO).(19) These cultures were infected with Cre-recombinase to delete all RIMBPs and RIMs, then rescued with either the RIMBP control (RIMBP-WT) or RIMBP mutant (RIMBP-MUT) (Figure 4B), and analyzed via patch-clamp or Ca^2+^ imaging (Figure 4C, D).

**Figure 4.**
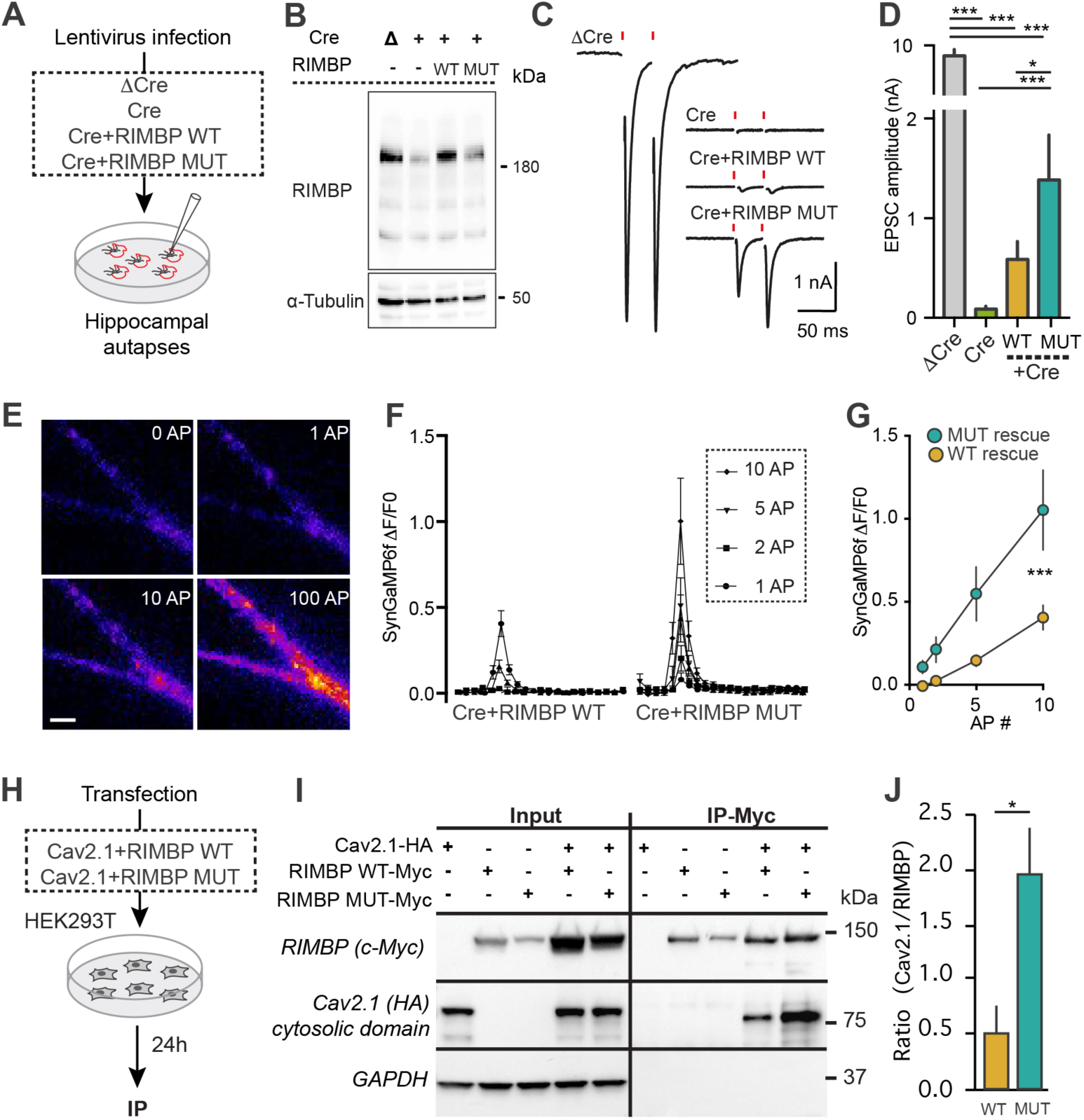
Impact of RIMBP1-p.Gly1808Ser variant on synaptic transmission. **A, B**. Hippocampal autapses for assessing the role of p.Gly1808Ser on synaptic transmission *in vitro*. **A**. Schematic of experimental configuration. Synaptic transmission in individual neurons making synapses onto themselves (autapses) was measured by whole-cell patch clamp recordings. Hippocampal autapses were prepared from RIMBP1,2/RIM1,2 quadruple conditional KO mice, and infected with lentiviruses expressing a recombinase-deficient version of Cre as a control (here after ‘ΔCre’), Cre-recombinase (‘Cre’), Cre+RIMBP WT, and Cre+RIMBP MUT constructs. **B**. RIMBP levels in the presence of lentiviruses expressing ΔCre, Cre, Cre+RIMBP WT, and Cre+RIMBP MUT. In this experiment, α-tubulin was used as a loading control. **C**. Representative recordings of single action-potential evoked release in hippocampal autapses. **D**. Summary graphs of single spike-evoked release in autapses recued with either RIMBP-WT or with the pathogenic RIMBP-MUT construct. **E-G**. Direct measurements of spike-triggered presynaptic calcium entry. **E**. Fluorescent images of a representative hippocampal dendrite expressing SynGCaMP6f under basal conditions (0 AP), or after 1, 10, and 100 action potentials. **F**. Time course of presynaptic fluorescence signals (ΔF/F0) as a function of time following 1, 2, 5, or 10 presynaptic action potentials. **G**. Summary graph of presynaptic fluorescence signals for different action potential frequencies in autaptic cultures infected with viruses expressing Cre+RIMBP WT and Cre+RIMBP MUT. **H-J**. RIMBP pathogenic missense variant impacts RIMBP-Ca^2+^-channel interaction. **H**. Experimental strategy for assessing the interaction of RIMBP with calcium-channel using co-immunoprecipitations (co-IPs). **I**. Co-immunoprecipitation experiments to test the interaction of RIMBP with Ca^2+^-channel -subunit. Cell lysates from HEK293T cells expressing CaV2.1-HA tagged and RIMBP WT-myc tagged, or CaV2.1-HA tagged and RIMBP MUT-myc tagged were subjected to immunoprecipitations with antibodies against myc. Input fractions (left; 1% of total) and immunoprecipitates (right, IP) were analyzed by immunoblotting with the following antibodies: myc- epitope (top) to recognize RIMBP, HA-epitope (middle) to identify CaV2.1, and GAPDH as a negative control (low). **J**. Summary graph of three co-immunoprecipitation experiments performed as in **I**. Number of experiments (cells/cultures): **C, D**. ΔCre (32/3); Cre (24/3); RIMBP WT (43/3), RIMBP MUT (35/3)**. F, G**. Cre+RIMBP WT (17/3), Cre+RIMBP MUT (16/3)**. J**. RIMBP WT (5), Cre+RIMBP MUT (5).

Deletion of all RIMBPs and RIMs completely blocked spike-triggered glutamate release from cultured neurons, as previously described (19) (Figure 4C, D). Surprisingly, neurons rescued with RIMBP-MUT showed, on average, nearly 100% larger excitatory postsynaptic current (EPSC) amplitude than RIMBP-WT, indicating that RIMBP1- p.Gly1808Ser variant does not compromise the function of RIMBP1 but rather causes abnormally increased synaptic transmission. As the pathogenic variant is located in the last SH3 domain of RIMBP1, which binds to VGCCs at the active zone and clusters them in close proximity of Ca^2+^sensors(17) (Figure 1D), we hypothesized that p.Gly1808Ser may affect release by regulating the interaction with VGCCs, which in turn would impact total Ca^2+^ entry during action potential firing. Thus, we directly measured intra-terminal Ca^2+^ levels using synaptically-localized SynGCamp6F in qKO autaptic cultures rescued with either RIMBP-WT or MUT (Figure 4E, J). Neurons transduced with the RIMBP-WT rescue construct showed moderate presynaptic Ca^2+^ elevations in response to increasing numbers of action potentials (Figure 4F, G). In striking contrast, neurons rescued with the pathogenic RIMBP-MUT variant displayed more than 100% larger Ca^2+^ transients (Figure 4G).

Finally, we studied whether p.Gly1808Ser affects the interaction of RIMBPs with VGCCs. We co-transfected HEK293T cells with a plasmid carrying an HA-tagged cytosolic C-terminal portion of the a1 subunits of P/Q (Cav2.1), known to mediate the interaction with RIMPBs, and a plasmid containing c-myc tagged RIMBP-WT or RIMBP-MUT (Figure 4H). Co-immunoprecipitation experiments revealed that RIMBP- MUT precipitated more than 3-fold higher amounts of Cav2.1 compared to RIMBP-WT (Figure 4I, J, p<0.03). Altogether, these experiments indicate that p.Gly1808Ser causes abnormally increased neurotransmitter release via a mechanism that involves enhanced spike-triggered presynaptic Ca^2+^ entry, likely by recruiting a higher number of VGCCs at the release sites.

## Discussion

We report here the identification of three pathogenic homozygous variants in *TSPOAP1*, the gene encoding the active zone protein RIMBP1, as a genetic cause of autosomal recessive dystonia. Two of these variants (p.Ala180Profs*8 and p.Gln817*) result in complete loss of RIMBP1 function either through early truncations and loss of all functional domains or through nonsense-mediated decay. The third variant (p.Gly1808Ser) is a missense variant located in the last SH3 domain of RIMBP1 and leads to abnormally increased synaptic transmission by likely recruiting more VGCCs to the active zone.

Clinically, the four subjects carrying homozygous truncating variants displayed severe and progressive form of generalized dystonia featuring prominent cranio-cervical involvement with onset in early teenage years. Strikingly, in these patients dystonia was also associated with pauci-symptomatic progressive cerebellar atrophy, a presentation that to date has been described in only a small number of families (31). On the other hand, the three subjects carrying the missense variant p.Gly1808Ser showed a milder phenotype featuring later onset focal dystonia, without clear evidence of significant cerebellar structural defects.

In mice, complete removal of RIMBP1 resulted in several motor abnormalities, including increased spontaneous locomotor activity, abnormal beam walking, and the presence of marked hindlimb clasping behaviors. Importantly, these motor behaviors are commonly observed across different genetic mouse models of dystonia (32), further supporting the pathogenic role of complete RIMBP1 loss-of-function in dystonia. Additionally, we were able to trigger dramatically abnormal movements and postures in response to subthreshold injections of the muscarinic agonist oxotremorine, known to unmask the appearance of dystonia-like movements in other genetic models of dystonia (26). Lastly, loss of RIMBP1 also resulted in structural abnormalities in the cerebellum, including shortening of the dendritic tree of Purkinje cells, reduced number of excitatory, but not inhibitory synapses, and abnormal cerebellar expression of Calbindin and Synapsin. Altogether, our results indicate that, like in humans, loss of RIMBP1 function in mice leads to progressive structural abnormalities in the cerebellum, more remarkably in cerebellar Purkinje cells, which might underlie, at least in part, the motor abnormalities we observed in RIMBP1-KO mice.

Our results indicate that RIMBP1, but not RIMBP2, is a key molecular link between synaptic dysfunction and dystonia. This is likely because in mice and humans RIMBP1 is heavily expressed in cerebellar Purkinje cells, where the homologous and functionally redundant RIMBP2 is virtually not expressed, thus preventing functional compensation and making Purkinje cells particularly susceptible to the effect of genetic variants affecting RIMBP1 function. Importantly, the association of cerebellar structural abnormalities and dystonia observed in both humans and mice lacking RIMBP1 supports the increasingly recognized role of cerebellar dysfunction in dystonia pathogenesis. For example, dystonia is increasingly recognized as one of the main presentation of several disorders in which pathology primarily involves the cerebellum, including ataxia-telangiectasia (33) and several autosomal dominant spinocerebellar ataxias (34). More mechanistically, recent work in mice showed that abnormal and irregular burst firing of Purkinje cells, as well as in neurons in the deep cerebellar nuclei, can be observed in several mouse models of dystonia (35-40). Furthermore, impaired cerebellar output may directly cause dystonia by altering basal ganglia activity and plasticity of cortico-striatal pathways (41). In this context, given the essential role of RIMBPs in regulating precise and reliable synaptic release (18), we speculate that RIMBP1 deletion may cause irregular synaptic transmission of Purkinje cells resulting in aberrant plasticity at cortico-striatal synapse. This effect is likely further precipitated by oxotremorine as over-activation of cholinergic M1 receptors on striatal neurons has been shown to have a strong effect on disrupting cortico-striatal synaptic plasticity (42).

RIMBPs are enriched at the interface between the active zone cytomatrix and the presynaptic membrane where they directly interact with RIMs and P- and Q-type VGCCs via their SH3 domains. Via these interactions, RIMBPs promote accumulation of VGCCs (i.e. Cav2.1 and Cav2.2) at the release sites, and thereby regulate the dynamics of transmitter release in response to presynaptic action potentials. Therefore, mechanistically, RIMBP1 variants can influence synaptic function by altering the clustering of release-relevant presynaptic VGCCs. Loss-of-function variants are expected to reduce the density of active zone VGCCs, as observed in central synapses of mice lacking RIMBPs (18, 19, 28, 43). Conversely, the missense variant found in family C (p.Gly1806Ser) increased the extent of transmitter in response to action potentials. We also observed that this variant increased presynaptic Ca^2+^ elevation per action potential and enhanced spike-triggered intraterminal Ca^2+^, likely due to an increased number of VGCCs per active zone. Together, this indicates that both decreased or increased density of VGCCs at the active zone is key in dystonia pathogenesis.

RIMBPs can also interact with presynaptic Ca^2+^ activated potassium channels (K-Ca^2+^ channels), in particular BK channels, via their central FN3-type domains and regulate their clustering at the release sites (44). Although the exact physiological significance of this interaction remains to be determined, it is likely that active zone BK channels limit the extent of membrane depolarization and transmitter release during sustained trains of presynaptic action potentials. Therefore, RIMBP1 variants may also alter the density of BK-channels. While our results indicate that the total amount of BK alpha subunit was not altered in RIMBP1-KO mice, it is not known if the nano-scale organization of BK channels might be impaired both in Purkinje cells as well as in other brain regions involved in dystonia pathogenesis. At the cellular level, K-Ca^2+^channel dysfunction leads to uncoordinated firing patterns in Purkinje cells, which is expected to alter the fidelity and reliability of information transfer to downstream targets in the deep cerebellar nuclei (45). This phenotype is identical to that observed upon genetic removal of RIMBPs from other central synapses (43).

Importantly, evidence from human genetic studies supports the role of perturbed presynaptic neurotransmitter release in the pathogenesis of hyperkinetic movement disorders. Pathogenic dominant variants in the RIMBP1 binding partners *CACNA1A* and *KCNMA1*, encoding Cav2.1 and BK-channels respectively, have been linked to a variety of human movement disorders, including episodic ataxia(46) and paroxysmal dyskinesias (47, 48). Moreover, recessive variants in *CACNA1A, CACNA1B* and *KCNMA1* cause progressive epileptic encephalopathy variably associated with cerebellar atrophy (49, 50) or dyskinesias (51). Furthermore, PNKD, another gene linked to a paroxysmal dyskinetic movement disorder, has been recently shown to localize at synapses where it interacts with RIM1 and RIM2 modulating neurotransmitter release (52). Finally, a gain-of-function missense variant in the presynaptic protein MUNC13A, causally linked to a complex dyskinetic movement disorder, was recently shown to cause pathologically enhanced neurotransmitter release (53), a synaptic phenotype similar to what we observed for the RIMBP1 variant p.Gly1806Ser.

In conclusion, our results show that RIMBP1 dysfunction causes dystonia in humans and abnormal movements and postures in mice. These results establish a causal link between presynaptic dysfunction and dystonia pathogenesis. As RIMBP1 dysfunction leads to unreliable synaptic transmission, we hypothesize that restoring the precision of synaptic release pharmacologically may represent a promising novel therapeutic angle to treat dystonia and other hyperkinetic movement disorders. The new RIMBP1- KO mouse model we developed could be instrumental not only to test this hypothesis, but also to disentangle the circuit abnormalities underlying dystonia pathogenesis and to further investigate the complex interaction between cerebellum and basal ganglia.

## EXPERIMENTAL PROCEDURES

### Patients

#### Gene mapping

DNA was extracted from peripheral lymphocytes following standard protocols. A genome-wide analysis genotyping scan was performed in all six members of family A and in the proband of family B using the HumanCytoSNP-12 DNA Analysis BeadChip Kit (Illumina, San Diego), according to manufacturer’s instruction. Homozygosity mapping was performed using PLINK v.1.079 (54). In family C, homozygosity mapping was performed using Whole-exome sequencing data analyzed with the H3M2 software (55).

#### Whole-exome sequencing

Whole-exome sequencing (WES) was performed in all three affected children and both parents for family A, in the proband only for family B and in the proband and her unaffected sibling for family C. WES and bioinformatic analysis were performed as previously described (56-58). Given the reported history of parental consanguinity in family A and the result of homozygosity mapping suggesting cryptic parental relatedness in family B and C, the analysis focused on homozygous coding and essential splice-site variants. Variants were not considered if they had a read depth < 5 and a minor allele frequency (MAF) > 0.005 in Exome Aggregation Consortium, Complete Genomics 69, 1000 Genomes project, and Exome Variant Server, as well as in in-house databases containing WES data from healthy controls or subjects with unrelated disorders. The summary of WES quality metrics for the families included in this study and full list of genetic variants surviving the initial filtering strategy are listed in supplemental tables 2 and 3.

### Mice

#### General

We used the following mouse lines: RIMBP1-KO, RIMBP2-KO, and quadruple RIM1,2/RIMBP1,2-KO. Mice were generated and maintained as described previously (18, 19).

#### Measurements of motor performance

All behavioral tests were performed in six-month-old mutant and control littermate male mice between 1 to 7 PM. 1. <u>Open field locomotion</u>. Mice were video-tracked in an open field arena for 60 minutes. Distance traveled, speed, and overall activity was analyzed using Viewer software. 2. <u>Limb-clasping.</u> Mice were suspended by their tails for 10, 20 or 30 seconds and the percentage of animals displaying clasping behavior at each time point was measured. To quantify limb-clasping, we used the following “clasping index”: ‘0’= no clasping, ‘1’ = single limb clasping, and ‘2’ =clasping in two or more limbs. In all experiments, mice were videotaped and analysis of video-recordings was performed offline in a blind-manner. 3. <u>Beam walking test.</u> Mice were trained to cross a suspended bean over 3 consecutive day and on the fourth day, they were video-taped and the time that took for them to cross the beam (10-90% of total distance) as well as the total number of ‘slips’ during the crossing, was quantified. 4.<u>Accelerating rotarod.</u> We used a five-station rotarod treadmill (ENV- 575M, Med Associates) equipped with an 8 to 80 rpm rate of acceleration over 300 s. Time and speed to fall off was quantified automatically (Med Associates). 5. <u>Motor behavior after systemic injections of oxotremorine</u>. Littermates wild type and RIMBP1-KO male mice were habituated in a clear plastic cage for 10 minutes and then treated with an intra-peritoneal injection of 0.01mg/kg of oxotremorine methiodide (Sigma-Aldrich) dissolved in normal saline solution (NaCl 0.9). Mice were videotaped for 60 minutes and motor behavior assessed every 10 minutes using a previously proposed scale for quantification of abnormal movements and postures (26). Two observers blinded to the genotype analyzed the experiment and rated the motor behavior of the mice.

#### Morphological measurements and synapse density

All morphological measurements were performed in six-month-old mice. To obtain a quantitative estimation of cerebellar size, we cut (50 mm) transversally PFA-fixed cerebelli, stained them with DAPI, imaged them under epifluorescence microscopy, and reconstructed them in 3D them using ImageJ. To assess Purkinje cell number, 40 um parasagittal sections from either RIMBP1 WT or RIMBP1-KO mice were immunostained with calbindin antibodies, imaged using confocal microscopy and analyzed with NIS-Elements AR software comparing the density of Purkinje cells by counting their somas. To measure excitatory synapse density, the cerebellum of 6 RIMBP1 WT and 6 RIMBP1 KO mice was cut parasagittal (30 um), and stained with antibodies against Calbindin (1:4000, mouse, Sigma) to label Purkinje cell dendritic profiles, and either vGluT2 (1:500, guinea pig, Millipore) or vGluT1 (1:1000, guinea pig, Millipore) antibodies to label climbing and parallel fiber input, respectively. To measure inhibitory synapse density onto Purkinje cells, 4 RIMBP1 WT and 4 RIMBP1 KO cerebelli were cut parasagittal (30 um) and stained with antibodies anti-GAD65 (1:1000, mouse, DSHB) and anti-calbindin (1:1000, rabbit, Millipore). Synaptic puncta were imaged using a Nikon confocal microscope (A1RSi+, Confocal Microscope, Nikon) controlled by the NIS-Elements AR software (Nikon). All the acquisition parameters were kept constant between wild-type and mutant conditions.

#### Neuronal cultures, electrophysiology, and calcium imaging

Autaptic cultures from RIM1,2/RIMBP1,2 KO (qKO) mice were prepared at P1. Briefly, neurons were plated on a sparse layer of glial micro-islands under growth-permissive substrate to obtain single neuron microislands. 24-36 h after plating, cultures were infected with either Cre-expressing or a recombination-deficient delta-cre-expressing vector as a control. For rescue experiments, neurons were additionally infected (24-36 hours post-plating) with either RIMBP-WT or with RIMBP-MUT constructs carrying the pathogenic p.G1808S variant. All autaptic cultures were grown for at least 2 weeks before recordings. Patch-clamp electrophysiology was performed at DIV 14-18 using electrode filled with an intracellular solution containing (in mM): 136 KCl, 17.8 HEPES, 1 EGTA, 4.6 MgCl_2_, 4 Na_2_ATP, 0.3 Na_2_GTP, and 12 creatine phosphate, and 50 U/ml phosphocreatine kinase (∼300 mOsm; pH 7.4). The extracellular solution contained (in mM): 140 NaCl, 2.4 KCl, 10 HEPES, 2 CaCl_2_, 4 MgCl_2_, and 10 glucose (pH adjusted to 7.3 with NaOH, ∼300 mOsm). Single action potentials were evoked via a 2 ms depolarization pulse (from -80 to 0 mV) in voltage-clamp while simultaneously recording evoked currents at -80 mV holding potentials.(30) For presynaptic Ca^2+^ imaging, autaptic cultures were additionally infected at DIV1 with a lentivirus expressing SynGCamp6. To evoke action potentials, neurons were patch-clamped as indicated above and then spikes were triggered by direct current injection in current-clamp. Images in Ca^2+^ imaging experiments were acquired using 490-nm LED system (pE2; CoolLED) at a 20 Hz sampling rate with a 50 ms exposure time.

#### Measurement of protein levels and co-immunoprecipitation

Protein levels were quantified in 6-month old mice via western blot. Briefly, cerebellar tissue was isolated manually, mechanically homogenized, solubilized in 2X Laemmli buffer, run in 4-20% Tris/Glycine gels, transferred onto nitrocellulose membrane, and immune-labeled with an array of specific antibodies again multiple synaptic and cerebellar markers (see full list in attached SOM). Membranes were analyzed with an Odyssey Infrared Imager.

For co-immunoprecipitations experiments, 5 × 10^6^ HEK293T cells were plated in 10-cm dishes and transfected with plasmids containing cDNA of mouse HA-tagged CaV2.1 (cytosolic C-terminal domain) and either rat myc-tagged RIMBP-WT or RIMBP-MUT (p.Gly1808Ser). The variant was introduced using Q5® Site-Directed Mutagenesis Kit (New England Biolabs). Cells were transfected with 6 μg of each plasmid using Lipofectamine 2000 (Invitrogen), harvested 24 hours after transfection. Cells were lysed on ice for 60 min lysed in EBC lysis buffer (Boston BioPRoducts) containing cOmplete™, Mini, EDTA-free Protease Inhibitor Cocktail (Roche, 11836170001) and subsequently centrifuged at 20,000 g for 10 min to remove cellular debris. Protein lysates were rotated overnight at 4 °C with anti-myc (Cell Signaling; 9B11), anti-HA antibodies (Cell Signaling; 6E2) or GAPDH (Millipore; MAB374). 25 µl of Dynabeads™ Protein G (Thermo Fisher Scientific) were then added to cell lysates and rotated at room temperature for 30 min. The samples were then washed 5X for 5 min at room temperature with lysis buffer. Dynabeads were finally resuspended in 2X Laemmli buffer after the final wash and analyzed by standard SDS/PAGE and immunoblotting.

#### Statistics

Data in figures 2, 3, and 4 are mean ± SEM. Statistical significance was assessed by ANOVA (Figure 2D, left and Figure 4D, G), Student’s t-test (Figure 2A, B, C, and D, right; Figure 3; Figure 4J). *p < 0.05, **p < 0.01, and ***p < 0.001; n.s., non-significant.

#### Study Approval

The genetic studies were approved by the local ethics committees (Family A - University College London Hospitals, London, UK; Family B - University of Tartu, Tartu, Estonia; Family C - Istanbul University, Istanbul, Turkey). Written informed consent for sample collection and subsequent analysis was obtained from all individuals or their guardians prior to inclusion in the study. All experiments involving mice were performed in accordance with Stanford and Federal Guidelines and were approved by the Stanford Institutional Animal Care and Use Committee and the animal and the animal ethics guidelines of the University of Heidelberg under license T-52/18 and G-268/18. Experiments involving hippocampal culture neurons were performed according to the regulations of Berlin authorities and animal welfare committee of the Charité–Universitätsmedizin Berlin, Germany under license no. 0220/09.

## Author contribution

Conception and design of the study: NEM, CA, TCS.

Funding acquisition: CA, DK, NWW, TCS, CR, KO, EL, TG

Conducting experiments: NEM, MMB, JD, SP, BA, PGL, CP, MS, GJC, JC, GP

Acquiring data: AT, AP, JSS, BB, SW, TTW, AP, AS, RR, LKE, SP, KR, TT, HH, KPB, MAK

Supervision: CA, DK, NWW, TCS, CR, KO, EL

Writing of the first draft of the manuscript: NEM, CA.

All authors contributed to revising the manuscript.

## Acknowledgments

NEM is funded by a Parkinson’s foundation training grant. MS received a Heisenberg Fellowship from the German Research Council (DFG). This work is supported by National Institute of Health R37 NS096241 (DK), the Chica and Heinz Schaller Stiftung, NARSAD Young Investigator, and DFG1158-Z01 (CA), the Estonian Research Council grants PUT355, PRG471 and PUTJD827 (SP, KO). We thank Dr Ülle Krikmann who provided some videos for patient B.

## Web Resources

The URLs for data presented herein are as follows:

1000 Genomes project: www.1000genomes.org

Allen Mouse Brain Atlas: http://mouse.brain-map.org/

CADD: http://cadd.gs.washington.edu/home

Clustal Omega: http://www.ebi.ac.uk/Tools/msa/clustalo/

Complete Genomics 69 database: www.completegenomics.com/public-data/69-Genomes

dbSNP: www.ncbi.nlm.nih.gov/projects/SNP

Exome Aggregation Consortium database: http://exac.broadinstitute.org/ (last accessed February 2019)

Genome Aggregation Database: https://gnomad.broadinstitute.org/ (last accessed February 2019)

GATK https://www.broadinstitute.org/gatk/

GeneMatcher https://genematcher.org/

Genotype-Tissue Expression (GTEx) project: https://gtexportal.org/home/

Great Middle East Variome database (http://igm.ucsd.edu/gme/index.php)

NHLBI Exome Variant Server EVS: evs.gs.washington.edu

Online Mendelian Inheritance in Man (OMIM), http://www.omim.org/

PolyPhen2: http://genetics.bwh.harvard.edu/pph2/

SIFT: http://sift.jcvi.org/

