## supplemental material for "Bi-allelic variants in *TSPOAP1*, encoding the active zone protein RIMBP1, cause autosomal recessive dystonia"

### Supplemental data

#### Supplemental Figures

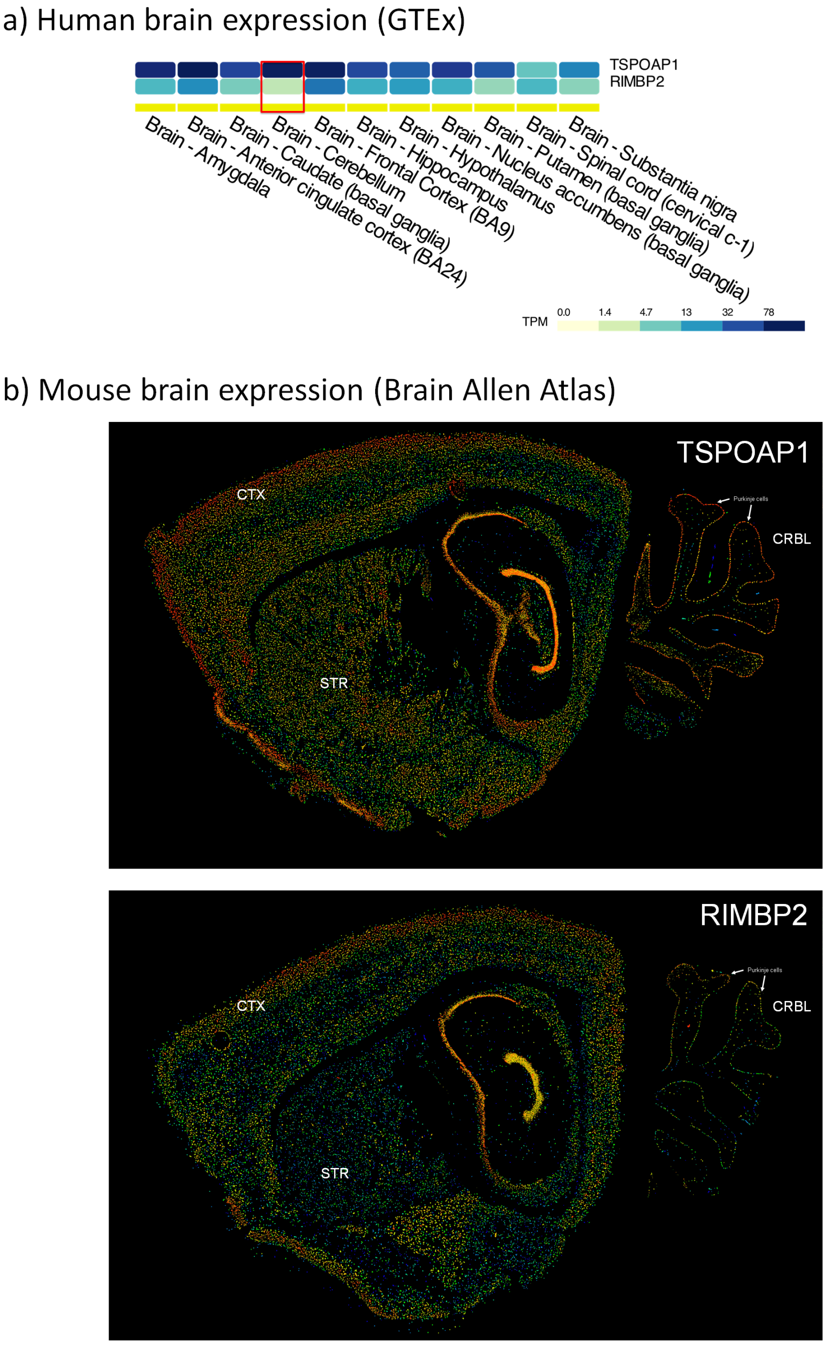


**Suppl. Figure 1. Expression patterns of RIMBPs in human and mouse brain.**
**A.** Expression in human brain. Levels of *TSPOAP1/*RIMBP1 and *RIMBP2* mRNA expression levels in eleven adult human brain regions (source: GTEx, see Web Resources). This dataset levels was generated with Illumina TrueSeq RNA sequencing and Affymetrix Human Gene 1.1 ST Expression Array (V3; 837 samples) and tissue originating from nearly 1000 donors. Gene and transcript expression levels on the GTEx Portal are shown in Transcripts Per Million (TPM).
**B.** Expression in mouse brain. *TSPAOPA1/* RIMBP1 and *RIMBP2* expression in the mouse brain in sagittal sections. Images were obtained from the Allen Mouse Brain Atlas (©2015 Allen Institute for Brain Science). Expression intensity is color-coded and ranges from low (blue) to moderate (green, yellow) to high (red) intensity.

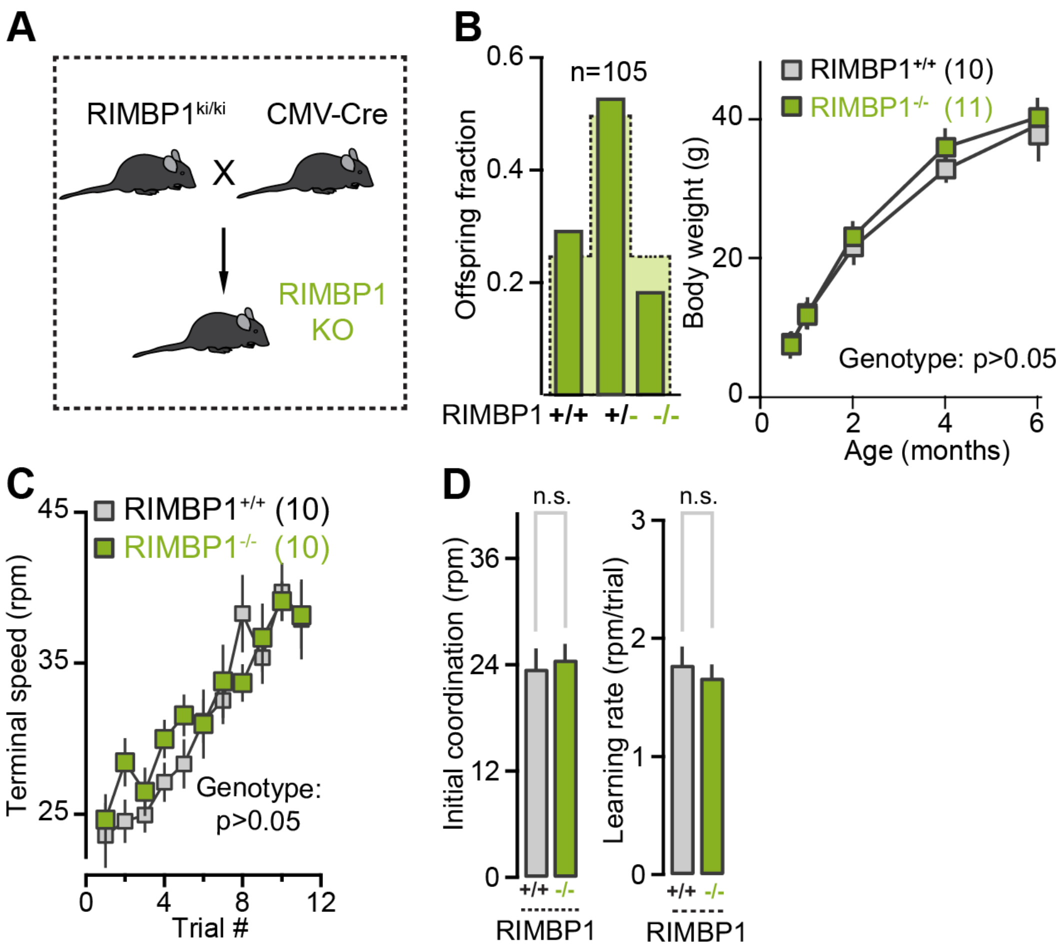


**Suppl. Figure 2. RIMBP1 constitutive knock-out mice.**

**A**. Schematic of crosses used to generate constitutive RIMBP1 KO and littermates control mice.

**B**. Left. Breeding offspring ratios resulting from RIMBP1 heterozygotes matings (105 mice were analyzed in total). Expected Mendelian ratios are shown by the dotted line. Right. Body weight of RIMBP1 WT and RIMBP1 KO mice.

**C.** Accelerating rota-rod test. Terminal speed as a function of trial number in RIMBP1 control and mutant mice.

**D**. Accelerating rotarod test. Left, summary graph of averaged initial motor coordination (terminal speed on day1, session1). Right, learning rate (slope of linear function from terminal speed vs trial # graph) in control and littermates KO mice.

Data are mean±SEM. Number of experiments: **C**, **D**. 10 WT, 10 KO. Statistical analysis was assessed by ANOVA (**C**) and Student's t-test (**D**); *$p < 0.05$, **$p < 0.01$, and ***$p < 0.001$; n.s., non-significant.

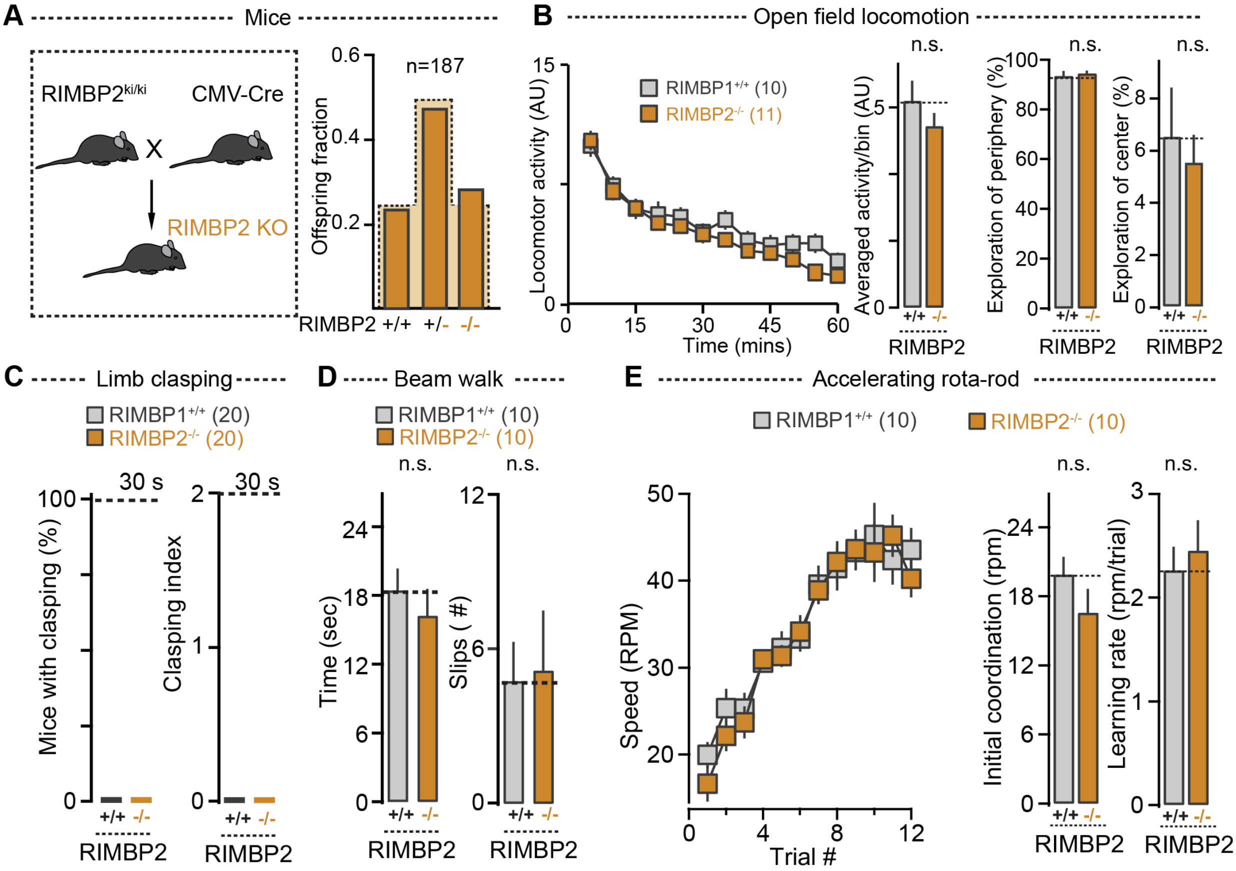


**Suppl. Figure3. Motor performance of RIMBP2 constitutive knock-out mice.**

**A**. Left, schematic of crosses used to generate constitutive RIMBP2 KO and littermates control mice. Right, breeding offspring ratios resulting from RIMBP2 heterozygotes matings (187 mice were analyzed in total). Expected Mendelian ratios are shown by the dotted line.

**B**. Open field locomotor activity. Left, locomotor activity as a function of time (bin width: 5 mins). Middle left, bar graph of total activity in RIMBP2 WT and RIMBP2 KO mice. Middle right, comparison of relative activity in the periphery (P) of the open field arena in control and mutant mice. Right, comparison of relative activity in the center (C) of the open field arena in control and mutant mice.

**C**. Limb-clasping test. Left, percentage of RIMBP2 WT and RIMBP2 KO mice displaying limb clasping in tail suspension test lasting 30 secs. Right, clasping index in control and mutant mice tail-suspended for 30 secs.

**D**. Beam-walk test. Left, summary graphs of total time to cross the beam. Right, summary graphs of total number of slips.

**E**. Accelerating rota-rod test. Left, terminal speed as a function of trial number in RIMBP2 control and mutant mice. Middle, summary graph of averaged initial motor coordination (terminal speed on day1, session1). Right, learning rate (slope of linear function from terminal speed vs trial # graph) in control and littermates KO mice.

Data are mean±SEM. Number of experiments: B, right: 10 WT, 11 KO; C: 20 WT, 20 KO; D: 10 WT, 10 KO; E: 10 WT, 10 KO. Statistical analysis was assessed by Student's t-test and ANOVA. $^{*}p < 0.05$, $^{**}p < 0.01$, and $^{***}p < 0.001$; n.s., non-significant.

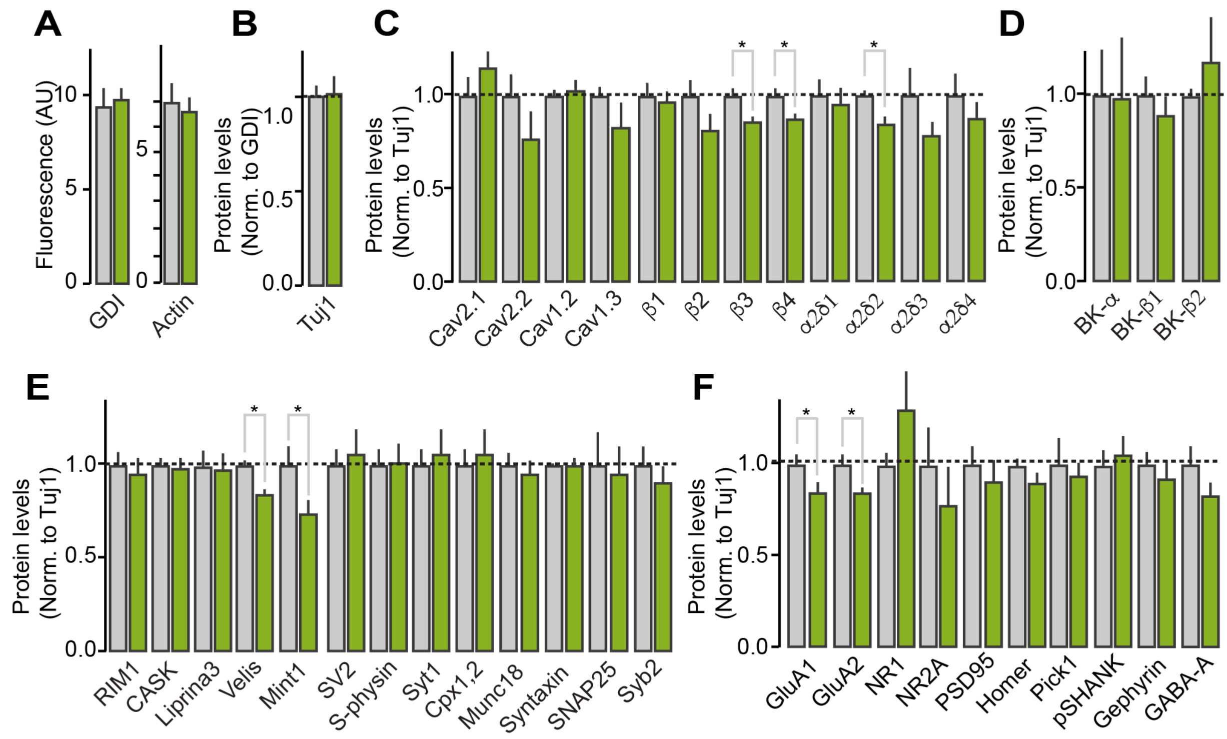


**Suppl. Figure 4. Biochemical composition of the cerebellum upon deletion of RIMBP1**.

**A.** Bar graphs summarizing the raw intensity of fluorescent signals arising from GDI and Actin after Western Blot analysis in RIMBP1 WT and KO cerebelli.

**B**. Summary graph showing Tuj1 signals normalized to GDI in RIMBP1 WT and KO cerebelli.

**C**. Summary graph of the levels of calcium-channel a-subunit, b-subunits, and a2d subunits in cerebelli from littermates RIMBP1 control and mutant mice.

**D**. Same as in **C** but BK-channels (a, b1, and b2).

**E**. Bar graphs showing the levels of several active zone, as well as of vesicle-associated proteins, SNAREs, and other presynaptic proteins that collaborate with SNAREs in synaptic vesicle fusion. All samples were obtained from cerebellar lysates from controls and RIMBP1 KO mice.

**F**. Summary graphs displaying the levels of several postsynaptic proteins and receptors for both excitatory and inhibitory neurotransmitters. Detailed information of the antibodies and dilution used can be found in the supplementary material.

Data are mean±SEM. Number of experiments (mice/sections): **A:** 12 WT, 12 KO, **B**: 10 WT, 10 KO, **C:** 7WT, 8 KO**;** **D**: 9 WT, 8 KO; **E**: 8 WT, 8 KO; **F:** 8 WT, 8 KO. Statistical analysis was assessed by Student's t-test. *p < 0.05 and ***p < 0.001; n.s., non-significant.

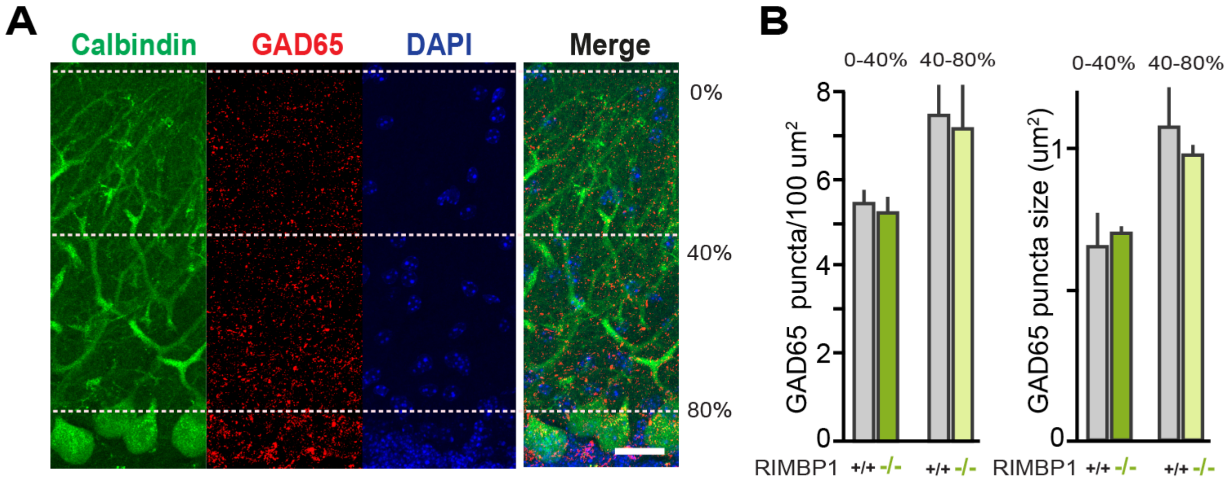


**Suppl. Figure 5. Cerebellar inhibitory synapses in RIMBP1 knock-out mice**.
**A.** Representative confocal images of cerebellar sections from a RIMBP1-WT mouse stained with antibodies anti-Calbindin (green, to label Purkinje cells), anti-GAD65 (red, to visualize inhibitory terminals), and with DAPI (blue, cell nuclei).
**B**. Summary graphs of GAD65 cluster density (puncta/100um$^2$, left) and GAD65 cluster size (right) in the outermost (0-40%) and innermost (40-80%) dendritic domains of Purkinje cells.

Data are mean±SEM. Number of experiments (mice/sections): 3/24 WT, 3/27 KO. Statistical analysis was assessed by Student's t-test. *p < 0.05 and ***p < 0.001; n.s., non-significant.

### Supplemental tables

#### Supplemental table 1. Standard quality metrics of whole-exome sequencing in families with *TSPOAP1* pathogenic variants

| Sample ID | MEAN_TARGET_COVERAGE (reads) | PCT_TARGET_BASES_2X (%) | PCT_TARGET_BASES_10X (%) | PCT_TARGET_BASES_20X (%) |
|---|---|---|---|---|
| A.II-1 | 61.78 | 98.79 | 95.53 | 87.64 |
| A.II-2 | 44.68 | 98.87 | 92.25 | 76.30 |
| A.II-3 | 44.65 | 98.29 | 86.43 | 67.82 |
| A.I-1 | 27.07 | 97.72 | 81.94 | 54.91 |
| A.I-2 | 98.04 | 99.23 | 97.67 | 94.34 |
| B.II-1 | 54.85 | 99.70 | 97.00 | 88.70 |
| C.II-2 | 82.02 | 98.08 | 95.82 | 90.84 |
| C.II-4 | 89.04 | 98.08 | 96.17 | 92.10 |

#### Supplemental table 2. Additional homozygous variants identified by whole-exome sequencing in families with *TSPOAP1* pathogenic variants

| Chr. Position | Ref | Alt | Gene | AA Change | gnomAD MAF | Homozyg./hemizyg. count in gnomAD | CADD | SIFT | PolyPhen-2 | Reasons to exclude pathogenicity |
|---|---|---|---|---|---|---|---|---|---|---|
| **Family A** | | | | | | | | | | |
| 14:20944705 | A | G | *PNP* | NM_000270:c.815A>G, p.E272G | 0.00006364 | 0 | 22.9 | T | B | Not in disease-linked ROH; unaffected father is homozygote; normal purine metabolism |
| 17:56403686 | C | - | *TSPOAP1* | NM_004758:c.538delG, p.A180fs | 0 | 0 | NA | NA | NA | |
| **Family B** | | | | | | | | | | |
| 14:45403616 | T | C | *KLHL28* | NM_017658:c.1045A>G, p.I349V | 0.003643 | 4 | 15.24 | T | B | Homozygous variants in gnomAD controls; in silico tools predict benign effect |
| 14:105418115 | C | T | *AHNAK2* | NM_138420:c.3673G>A, p.G1225S | 0.0002529 | 18 | 10.13 | T | B | Homozygous variants in gnomAD controls; in silico tools predict benign effect |
| 17:56389733 | G | A | *TSPOAP1* | NM_004758:c.2449C>T, p.Q817X | 0 | 0 | 35 | NA | NA | |
| 17:56436023 | G | A | *RNF43* | NM_017763:c.1114C> T, p.P372S | 0.000234 | 0 | 10.13 | T | B | In silico tools predict benign effect |
| 17:59489823 | C | A | *C17orf82* | NM_203425:c.487C>A, p.Q163K | 0 | 0 | 14.42 | T | D | In silico tools predict benign effect |
| X:101139206 | T | C | *ZMAT1* | NM_001282400:c.680A>G, p.D227G | 0.000009771 | 0 | 25.6 | D | D | |
| X:135630910 | G | A | *VGLL1* | NM_016267:c.377G>A, p.R126Q | 0.0006238 | 55 | 2.726 | T | B | Hemizygous variants in gnomAD controls; in silico tools predict benign effect |
| **Family C** | | | | | | | | | | |
| 8:36692371 | T | C | *KCNU1* | NM_001031836:c.1280T>C, p.I427T | 0.0001083 | 0 | 8.867 | T | B | Affected siblings are both heterozygous; in silico tools predict benign effect |
| 11:71174492 | G | A | *NADSYN1* | NM_018161:c.278G>A, p.R93Q | 0.00002123 | 0 | 23.1 | T | B | Affected siblings are heterozygous (C-II.1) and wild-type (C-II.3) |
| 17:56382759 | C | T | *TSPOAP1* | NM_004758:c.5422G>A, p.G1808S | 0.00001061 | 0 | 27.7 | D | D | |

Ref=reference allele; Alt=alternative allele; AA=amino acid; MAF=minor allele frequency; T=tolerated; B=benign; D=deleterious/damaging

#### Supplemental Movies

**Video 1. Family A and Family B.** Subject A-II.2. Exam shows severe dysarthria with mild orofacial dystonic movements with speech. There is elevation of the right shoulder and cervical dystonia with mild torticollis to the right. There is continuous dystonic posturing of bilateral upper extremities with finger flexion, wrist flexion and arm supination, particularly of left arm. Dystonic posturing worsens and there is muscle overflow when subjects is asked to hold arms extended in front of him as well as when attempting cerebellar maneuvers. There is no bradykinesia, but finger tapping movements are impaired by dystonic posturing of hands and there is also overflow to other forearm and arm muscles. Gait is narrow based, with bilateral foot inversion, worse on the right foot and dystonic posturing of bilateral upper extremities. Subject A-II.4. also shows whispering dysphonia suggestive of severe laryngeal dystonia. With speech, there are dystonic orofacial movements, also dystonic lingual movements. There is also left shoulder elevation and continuous dystonic posturing of right arm. Eye movement exam shows difficulty initiating saccades in all directions. There is bilateral dystonic posturing of upper extremities which worsen in severity when asked to hold arms extended in front of him and with cerebellar testing. Gait is narrow based, there is poor arm swing with dystonic posturing of right upper extremity with arm slightly abducted, extended wrist and fingers making a fist; there is slight foot eversion bilaterally and subject drags both feet when walking. The third segment shows subject B-II.1. At 17 years old there is generalized dystonia involving orofacial muscles, as well as upper extremities, with right arm abducted and held at the back and continuous dystonic posturing of left arm when attempting to use it, fingers are flexed and form a fist. Gait is narrow based, there is bilateral foot inversion, more prominent on the left, also with left leg circumduction; upper extremities are held away from his body with both hands making a fist. There cervical dystonia with anterocaput which is most noticeable towards the end of this segment. At 23 years old, gait has worsened, there is right foot inversion and knee flexion, both arms are held away from his body with hands making a fist, there is torticollis to the right, orofacial dystonia and blepharospasm. Orofacial and mandibular dystonia make tasks such as drinking or eating difficult, which is showcased in the last two segments.

**Video 2. Family C.** Subject C-II.4. On exam there is right shoulder elevation, mild right laterocollis, slight torticollis to the left and with slight retrocollis. Range of motion is full in all directions. Gait presents no abnormalities. Subject C-II.3. Examination shows a "no-no" head dystonic tremor. There is a mild, low frequency, high-amplitude postural tremor with upper limbs outstretched, most noticeable on index fingers. There is no evidence of dysarthria or bradykinesia. Gait shows good velocity and amplitude with slightly decreased arm swing bilaterally.

**Video 3. Beam walking test in *TSPOAP1*/RIMBP1 wild-type (WT) and knock-out (KO) mice.** Typical examples of beam crossing of a RIMBP-WT and a RIMBP1-KO mouse, showing a striking differences between genotypes in both the time to cross the beam as well as the total number of slips.

**Video 4 Hindlimb clasping in *TSPOAP1*/RIMBP1 wild-type (WT) and knock-out (KO) mice.** Side-by-side display of tail suspension test used to assess animal limb clasping.
Hindlimb clasping behaviour was consistently observed in RIMBP1-KO but not in RIMBP1-WT mice.

**Video 5. Motor behavior comparison between *TSPOAP1*/RIMBP1 knock-out (KO) mice and TSPOAP1/RIMBP1 wild-type (WT).** First segment shows motor behavior at baseline; RIMBP1-KO presents with mildly ataxic gait compared to RIMBP1-WT. Second segment shows behavior 20 minutes after oxotremorine 0.01 mg/kg intraperitoneally. RIMBP1-KO displays severely limited ambulation with splayed posture and intermittent body and head jerks. At 60 minutes after oxotremorine injection, RIMBP1-KO still displays slowed movement with hindlimbs infrequently in a splayed posture.

**Video 6. Examples of abnormal postures and movements in *TSPOAP1*/RIMBP1 knock-out (KO) mice.** First segment shows a mouse with moderate impairment, with repetitive jerking of snout, limited ambulation, splayed posture and brief forelimb movements resembling the initial phases of grooming. The second segment shows severe impairment with a hunched posture and little movement, with frequent head jerks. The third segment shows a mouse with severe impairment, with limited movement and a sustained single raised paw.

#### List of antibodies used for Western Blot analysis

| Antigen | Antibody | Dilution |
|---|---|---|
| RBP-1 (RIM-Binding Protein-1) | 316003 | 1:1000 |
| RBP-2 (RIM-Binding Protein-2) | 4193 | 1:1000 |
| RIM 1 (RIM1 Central Domains) | R809 | 1:2000 |
| RIM 2 (RIM2 C-Terminus) | T2795 | 1:250 |
| Liprin α3 | 4396 | 1:5000 |
| CASK | 75-000 | 1:1000 |
| Mint1 | P932 | 1;1000 |
| Veli1,2,3 | T813 | 1:1000 |
| Syntaxin 1 | HPC-1 | 1:1000 |
| SNAP 25 | 71.1 | 1:2000 |
| Synaptobrevin 2 | 69.1 | 1:2500 |
| Munc 18.1 | K329 | 1:500 |
| Complexin 1/2 | L669 | 1:1000 |
| Synaptotagmin 1 | 41.1 | 1:500 |
| Synaptophysin | 7.2 | 1:5000 |
| SV 2 | U1129 | 1:1000 |
| Synapsin1,2 | E28 | 1:500 |
| Calbindin | CB-955 | 1:1000 |
| GluA1 | 182003 | 1:1000 |
| GluA2 | 182103 | 1:1000 |
| NR1 | 75-272 | 1:1000 |
| NR2A | 75-288 | 1:1000 |
| PSD95 | 73-028 | 1:1000 |
| HOMER | 160003 | 1:1000 |
| PICK1 | U58317 | 1:1000 |
| SHANK | 75-089 | 1:1000 |
| Gephyrin | 147111 | 1:1000 |
| GABA-A | 224211 | 1:1000 |

| | | |
|---|---|---|
| $Ca^{2+}$v1.2-α1C Voltage-Gated Channel | ACC-003 | 1:500 |
| $Ca^{2+}$v1.3-α1D Voltage-Gated Channel | ACC-005 | 1:500 |
| $Ca^{2+}$v2.1-α1A Voltage-Gated Channel | ACC-001 | 1:500 |
| $Ca^{2+}$v2.2-α1B Voltage-Gated Channel | ACC-002 | 1:500 |
| $Ca^{2+}$β1 Voltage-Gated Channel | ACC-106 | 1:500 |
| $Ca^{2+}$β2 Voltage-Gated Channel | ACC-105 | 1:500 |
| $Ca^{2+}$β3 Voltage-Gated Channel | ACC-008 | 1:500 |
| $Ca^{2+}$β4 Voltage-Gated Channel | 75-054 | 1:500 |
| $Ca^{2+}$v-α2δ1 Voltage-Gated Channel | ACC-015 | 1:500 |
| $Ca^{2+}$v-α2δ2 Voltage-Gated Channel | ACC-102 | 1:500 |
| $Ca^{2+}$v-α2δ3 Voltage-Gated Channel | ACC-103 | 1:500 |
| $Ca^{2+}$v-α2δ4 Voltage-Gated Channel | ACC-104 | 1:500 |
| $BK_{Ca}{}^{2+}$- α | APC-021 | 1:500 |
| $BK_{Ca}{}^{2+}$- $\beta_1$ | APC-036 | 1:500 |
| $BK_{Ca}{}^{2+}$- $\beta_2$ | APC-034 | 1:500 |
| TUJ1 | T2200 | 1:4000 |
| GDI | 81.2 | 1:2000 |
| NeuN | ABN78 | 1:1000 |
